# iCn3D, a Web-based 3D Viewer for Sharing 1D/2D/3D Representations of Biomolecular Structures

**DOI:** 10.1101/501692

**Authors:** Jiyao Wang, Philippe Youkharibache, Dachuan Zhang, Christopher J. Lanczycki, Renata C. Geer, Thomas Madej, Lon Phan, Minghong Ward, Shennan Lu, Gabriele H. Marchler, Yanli Wang, Stephen H. Bryant, Lewis Y. Geer, Aron Marchler-Bauer

## Abstract

**Summary:** iCn3D (I-see-in-3D) is a web-based 3D molecular structure viewer focusing on the interactive structural analysis. It can simultaneously show 3D structure, 2D molecular contacts, and 1D protein and nucleotide sequences through an integrated sequence/annotation browser. Pre-defined and arbitrary molecular features can be selected in any of the 1D/2D/3D windows as sets of residues and these selections are synchronized dynamically in all displays. Biological annotations such as protein domains, single nucleotide variations, etc. can be shown as tracks in the 1D sequence/annotation browser. These customized displays can be shared with colleagues or publishers via a simple URL. iCn3D can display structure-structure alignment obtained from NCBI’s VAST+ service. It can also display the alignment of a sequence with a structure as identified by BLAST, and thus relate 3D structure to a large fraction of all known proteins. iCn3D can also display electron density maps or electron microscopy (EM) density maps, and export files for 3D printing. The following example URL exemplifies some of the 1D/2D/3D representations: https://www.ncbi.nlm.nih.gov/Structure/icn3d/full.html?mmdbid=1TUP&showanno=1&show2d=1&showsets=1.

**Availability and implementation:** iCn3D is freely available to the public. Its source code is available at https://github.com/ncbi/icn3d.

**Supplementary information:** User instructions are available at Bioinformatics online

## 1 Introduction

With the widespread availability of powerful personal computers and mobile devices, the visualization of biomolecular 3D structure (Kozlikova, et al., 2017; Sali, et al., 2015) is no longer restricted to stand-alone 3D viewer applications such as Chimera (Goddard, et al., 2018; Pettersen, et al., 2004), PyMol (Schrödinger, 2015), or Cn3D (Wang, et al., 2000), which all require the installation of a dedicated application. Interactive visualization of large molecular structures can now be achieved via web-based 3D viewers such as NGL (Rose, et al., 2018; Rose and Hildebrand, 2015), LiteMol (Sehnal, et al., 2017), or Aquaria (O’Donoghue, et al., 2015). Besides 3D visualization, 3D viewers also work as analysis tools to relate structures to functions and annotations. Table 1 summarizes features of some WebGL-based and stand-alone 3D viewers. A more complete list of 3D viewers can be found at https://en.wikipedia.org/wiki/List_of_molecular_graphics_systems, among many other listings.

**Table 1.** Feature comparison of some 3D viewers.

| Name | Web-based | 1D Sequence | 2D Diagram | Annotation | Align<sup>a</sup> | Share Link | EM Density Map | 3D Printing | Virtual Reality |
| --- | --- | --- | --- | --- | --- | --- | --- | --- | --- |
| iCn3D | ? | ? | ? | ? | ? | ? | ? | ? |  |
| LiteMol | ? | Web<sup>b</sup> | Web<sup>b</sup> | Web<sup>b</sup> |  |  | ? |  |  |
| NGL | ? | Web<sup>b</sup> |  | Web<sup>b</sup> |  |  |  |  |  |
| Aquaria | ? | ? |  | ? |  |  |  |  |  |
| Chimera |  | ? |  | ? |  |  |  | ? | ? |
| PyMol |  | ? |  | ? |  |  |  |  |  |
| Cn3D |  | ? | Web<sup>b</sup> | ? | ? |  |  |  |  |

<sup>a</sup>Align a protein sequence with unknown structure to a 3D structure.
<sup>b</sup>The feature is only available in a web page embedded with the viewer.

Previously, NCBI has developed and maintained “Cn3D” (Wang, et al., 2000), a stand-alone application written in C++. Taking advantage of advanced web technology, we have developed WebGL-based 3D viewer iCn3D, which covers many of the features provided by Cn3D, such as sequence displays in sync with the 3D structure displays. iCn3D also introduces novel features as suggested in “Twelve Elements of Visualization and Analysis” (Youkharibache, 2017). As described below, these features mainly include:

1. “Share Link” URLs to share visual displays of interest over the internet, with all the underlying annotations and user-provided labels;
2. selection behavior synchronized between displays of 3D structure, 2D interaction schematic, 1D sequence, and a menu of (pre-)defined sets, many of which are defined automatically during an interactive session to track user selections;
3. annotations such as the location of non-synonymous single nucleotide polymorphisms (SNPs) that have been automatically extracted and mapped to 3D structures from NCBI dbSNP database (Sherry, et al., 2001), protein domains, functional sites, molecular contacts, disulfide bonds, and custom annotations that can be defined interactively or through file/text based mechanisms, and can be displayed as “custom tracks” in the sequence/annotation browser;
4. visualization of sequence-structure alignment; and
5. visualization of structure-structure alignment and corresponding superposition.

## 2 Features

### 2.1 Share Link

The “Share Link” feature in iCn3D generates a URL that captures the commands used to customize the display of a structure. The URL can then be shared with others, enabling them to reproduce the same display. For example, Figure 1A shows a custom display of tumor suppressor P53 interacting with DNA (PDB ID 1TUP). iCn3D highlights key residues involved in the interaction: arginine 175, 248-249, 273 and 282, and glycine 245 (Cho, et al., 1994). The residue Arg 248 at chain B inserts into the major groove of the DNA double-helix. Other highlighted residues are also involved in protein-DNA interactions or protein-protein interactions. Mutations of these key residues may affect the interaction of P53 with DNAs and are associated with cancer (Vogelstein, et al., 2000). These mutations are shown on ClinVar (variants associated with clinical diseases) (Landrum, et al., 2018) and SNP sequence annotation tracks (Figure 1D).

**Figure 1:**
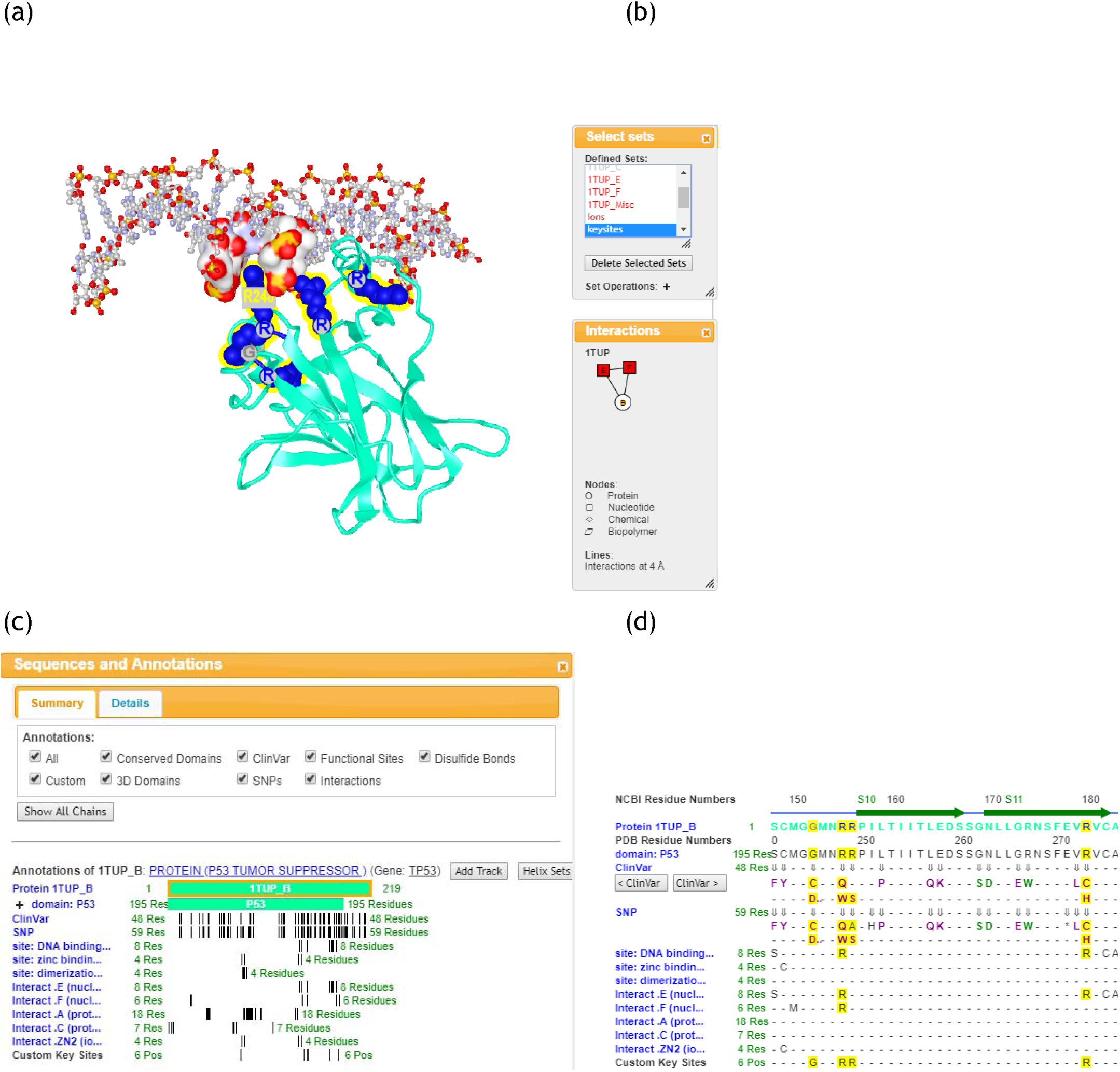
iCn3D visualizing the structure of tumor suppressor P53 complexed with DNA (PDB code 1TUP). This display can be reproduced in a live browser window with the shortened sharable link: https://icn3d.page.link/2rZWsy1LZmtTS3kBA. (a) “iCn3D PNG Image” saved using the option “Save File” in the File menu. (b) A list of defined sets including chains and custom sets (top), and a 2D schematic of the molecules involved in the complex structure (bottom). (c) Summary view of sequences and annotations with residues represented as vertical bars. (d) Detailed view of sequences and annotations.

iCn3D has made it easy to save a custom display and share it with others. Instead of saving the state of all atoms, iCn3D saves a command for each step. Like most 3D viewers, iCn3D can save a state file containing the command history. This state file can be used to reproduce the custom display. Since iCn3D is web-based, it further simplified the state file and concatenated commands in a single sharable link using the option “Share Link” in the File menu. The Share Link can be shortened, e.g. to https://icn3d.page.link/2rZWsy1LZmtTS3kBA, as shown above.

To save a custom display, it is recommended to save the image using the option “Save File > iCn3D PNG Image” in the File menu. This PNG image is of high quality, has a transparent background, and contains the sharable link at the end of the file. It can be imported to reproduce the display using the option “Open File > iCn3D PNG Image” in the File menu. Thus, this saved iCn3D PNG Image contains both the static image display and the commands to reproduce the dynamic display.

### 2.2 Synchronized Selection among 3D structure, 2D Interactions, Defined Sets, and 1D Sequences and Annotations

2D or 1D representations often convey information that is difficult to spot or highlight in complex 3D representations (Goodsell and Jenkinson, 2018; Heinrich, et al., 2014; Munzner, 2014). To this end, iCn3D visualizes macromolecular complexes in 3D, 2D and 1D, and synchronizes the selection among the various representations. The Share Link https://icn3d.page.link/2rZWsy1LZmtTS3kBA shows, in a single page, the 3D display (Figure 1A), 2D interaction schematic (Figure 1B bottom), 1D sequences (Figure 1D), and predefined sets including individual molecules/chains and custom sets (Figure 1B top). At the moment, iCn3D cannot show 2D topology diagrams. It provides two alternative displays instead. The first display is a new style called “Schematic” for proteins and nucleotides. This style shows the residues with circular 2D labels and connects the residues with 3D cylinders, as a combination of both 2D and 3D visualization styles. The second display is the 2D interaction schematic as shown in the bottom of Figure 1B. This display shows each protein/nucleotide/chemical as a node and shows interactions between them as lines. iCn3D presents 1D sequence in two different ways. Figure 1D shows the detailed sequence. Figure 1C shows the summary of the sequence and displays individual residue positions as vertical bars.

To simplify 3D views, iCn3D can display a selected subset only, using the option “View Only Selection” in the View menu. Figure 1A shows only the chain B interacting with DNA chains E and F while the protein chains A and C are not shown. Aside from displaying/hiding selections, iCn3D can also highlight selected sets of molecules and/or residues, or apply specific display style and/or color settings to a selection. The overall “Selection” mode, which can be toggled with a switch next to the Help menu, is very handy for command-based operations, determining whether future commands will apply only to the current selection or to the whole structure that has been loaded. iCn3D supports picking. Users can choose a selection option in “Select on 3D” in the Select menu, hold the “Alt” key, and select an Atom, Residue, Strand/Helix, or Chain in the 3D display (Figure 1A). Users can also click on the sets in “Defined Sets” (Figure 1B top) to select a set; click on the nodes or lines in 2D interaction schematic (Figure 1B bottom) to select a molecule or residues contacting that molecule; click on a title of a track in “Sequences and Annotations” (Figures 1C and 1D) to select an annotation; or click-and-drag on the sequence display to select a range of residues. An “Advanced” selection option using detailed specifications is also available in the Select menu.

The selections in the 3D, 2D and 1D displays are synchronized. As shown in Figure 1A, the key residues involved in the interaction of P53 with DNA are highlighted. This set is selected in “Defined Sets” (Figure 1B top), highlighted in the node B in 2D interaction schematic (Figure 1B bottom), and highlighted in the sequence (Figure 1D). Meanwhile, only the sequence of chain B is shown in the “Sequences and Annotations” window to simplify the user interface.

### 2.3 Annotations and Custom Tracks

Annotations are useful in the analysis of sequence-structure-function relationships. iCn3D provides several annotations originating from various NCBI databases: SNPs (from dbSNP (Sherry, et al., 2001)), ClinVar (annotated SNPs from ClinVar (Landrum, et al., 2018)), Conserved Domains (from CDD (Marchler-Bauer, et al., 2017)), 3D Domains (from MMDB (Madej, et al., 2014)), Functional Sites (from CDD), Interactions (from MMDB), and Disulfide Bonds (from MMDB). Like many genome browsers (Agarwala, et al., 2018; Kent, et al., 2002; Tang, et al., 2018), iCn3D lists annotations as tracks. Instead of using multiple level of zooming in and zooming out, iCn3D uses two levels of views: a “Summary” view with all residues fit in the available window (Figure 1C) and a “Details” view with the sequence shown as individual residues (Figure 1D). As shown in Figure 1D, Residues R248 and R273 appear on several tracks: domain P53, ClinVar, SNP, DNA binding site, Interacting with nucleotide chains, and the custom track “Custom Key Sites”. The comparison among these annotations show that R248 and R273 are directly interacting with DNA, and that their mutations have been characterized as pathogenic, as shown in the tooltip of residues on the ClinVar or SNP track.

Users may have their own annotations and may want to compare them with other annotations. iCn3D has several ways to add custom tracks, via NCBI Accession, FASTA sequence, BED File, Custom text, or Current Selection. Figure 1D shows the custom track “Custom Key Sites” added with the option of “Current Selection”.

### 2.4 Visualization of Sequence-Structure Alignment

Proteins with experimentally determined 3D structures represent only a small fraction of all known protein sequences. To relate 3D structure information to other proteins with similar sequences, homology models are often built for these proteins with unknown structures. Alternatively, iCn3D can show a sequence alignment of a query protein (e.g. accession NP_001108451.1) to a structure-derived sequence (e.g. accession 1TSR_A), together with the 3D coordinates and annotations available for that structure (Figure 2). The sequence alignment is highlighted in a box with three lines: the structure-derive sequence (e.g. “Protein 1TSR_A”), alignment details including matching residues (e.g. “BLAST E: 1.5e-84”), and the sequence (e.g. “Query: NP_001108451.1”). The 3D structure 1TSR_A in Figure 2 was colored by sequence conservation, which is calculated based on the BLOSUM62 matrix as used by BLAST (Altschul, et al., 1997; Altschul, et al., 2005). Red and blue colors indicate conservative and non-conservative substitutions, respectively. The 3D information and the annotations of the structure may provide useful hints with respect to the structure and function of the query protein.

**Figure 2:**
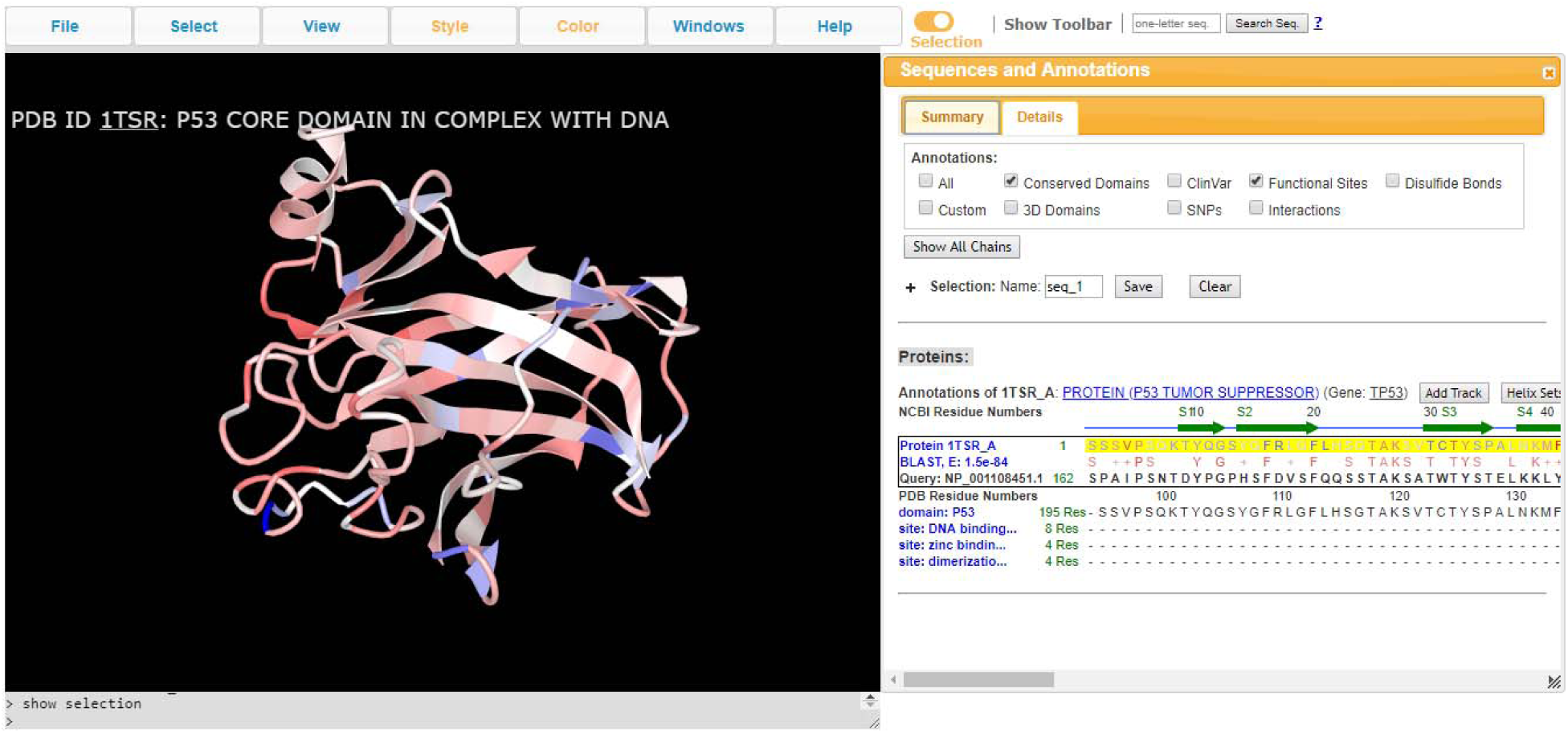
iCn3D visualizing the sequence alignment of protein accession NP_001108451.1 with structure accession 1TSR_A (chain A of 3D structure 1TSR). The alignment is shown in the box around “Protein 1TSR_A” in the sequence window. (Share Link: https://www.ncbi.nlm.nih.gov/Structure/icn3d/full.html?from=icn3d&blast_rep_id=1TSR_A&query_id=NP_001108451.1)

Users have two ways to align sequence to structure. First, click the “Align > Sequence to Structure” option in the File menu and input both a sequence ID and a structure ID. Second, click “Add Track” in the Sequences and Annotations window next to the chain “1TSR_A” and choose “NCBI gi/Accession”.

### 2.5 Visualization of Structure-Structure Alignment

iCn3D enables users to visualize structural alignments and superpositions for similar protein complexes. Structure alignments are pre-calculated by NCBI’s VAST+ service (Madej, et al., 2014). As shown in Figure 3, the 3D structure of oxy-hemoglobin (1HHO) is aligned with that of deoxy-hemoglobin (4N7N). All matching chains are superposed. Users can alternate between these two structures by pressing the key “a”. The aligned residues are colored in red for identical residues or blue for non-identical residues. In the 2D interactions window (Figure 3 middle), pairs of aligned protein molecules have the same number in red, e.g., both nodes 1HHO_A_1 and 4N7N_C are labeled with the number 3 colored in red.

**Figure 3:**
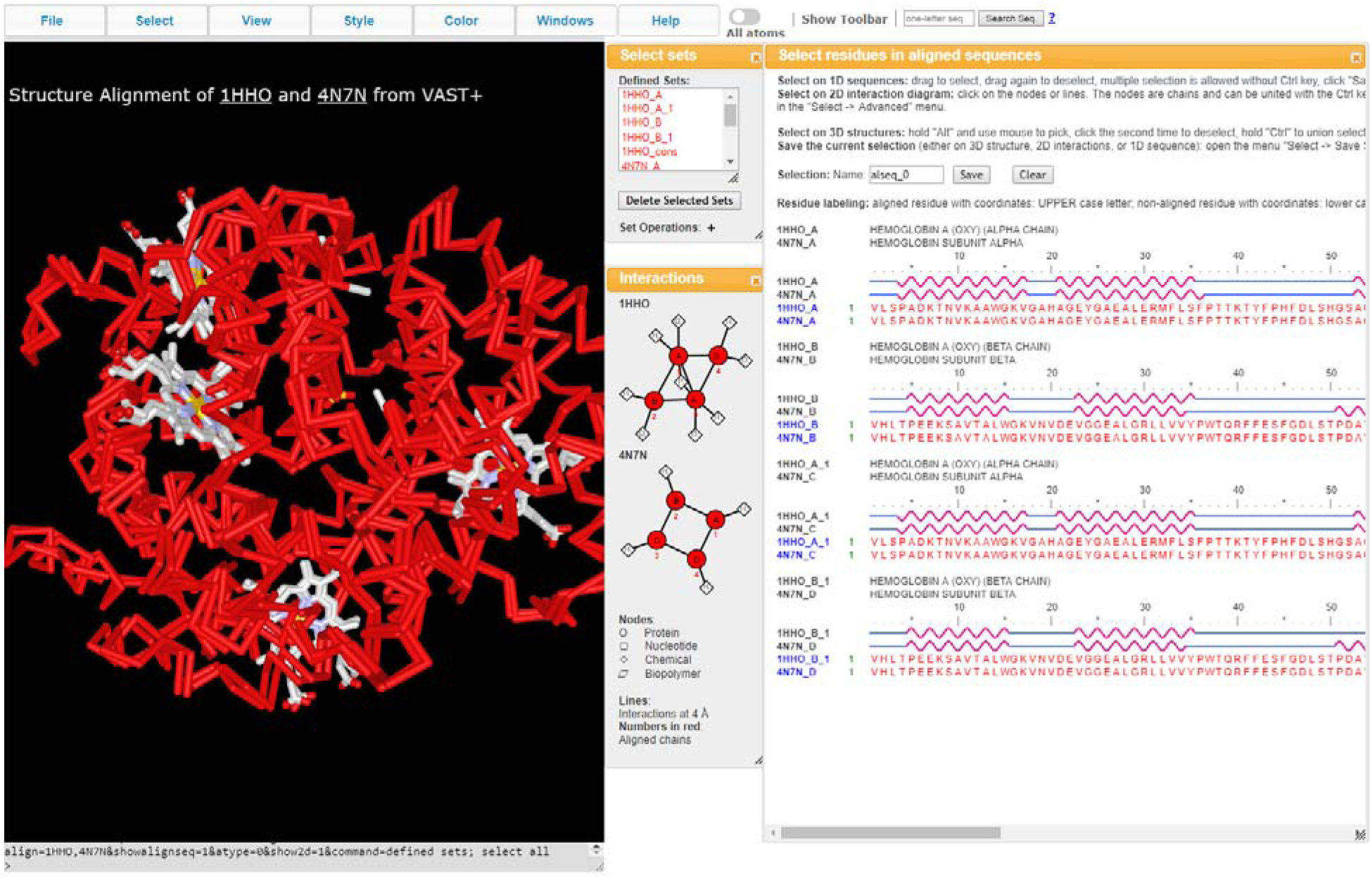
iCn3D visualizing the structure superposition and corresponding alignments between human oxy and deoxyhemoglobin (PDB ID 1HHO and 4N7N), as computed by VAST+. (Share Link: https://icn3d.page.link/wPoW56e8QnzVfuZw6)

## 3 Discussion

Third-party annotations, such as mutations from ClinVar or dbSNP, functional sites, interaction interfaces, structural or conserved domains, and their simultaneous visualization in 1D/2D/3D can provide useful and compelling evidence relating sequence, structure and function. More detailed genome level annotations could be linked to the structures in the future. For example, protein sequences could be shown together with chromosome, gene, intron, and exon in a genome browser such as https://www.ncbi.nlm.nih.gov/genome/gdv/browser/?context=gene&acc=7157.

iCn3D follows the FAIR (Findable, Accessible, Interoperable, and Reusable) guiding principles (Mons, et al., 2017; Wilkinson, et al., 2017). The JavaScript code of iCn3D is componentized to be reusable. The annotations in iCn3D could also be retrieved by other tools. In the future, iCn3D may adopt emerging 3D technologies such as virtual reality (Goddard, et al., 2018; Goddard, et al., 2018; Olson, 2018).

## Supporting information

User instructions

## Acknowledgments

We would like to thank Dr. Terry S. Yoo and Dr. David T. Chen for their helpful discussions on generating models for 3D printing. Dr. Alexander S. Rose provided valuable suggestions on reading MMTF files and implementing impostor methods. Chris Maloney helped to set up the building script using gulp. We also thank Qiangling Li, Spencer Bliven, Eli Draizen, Jose Duarte, Keiichiro Oto, and Tim Schaefer for their hackathon on the 2D interaction schematic during the 2016 ISMB meeting.

