## Supplementary material for "iCn3D, a Web-based 3D Viewer for Sharing 1D/2D/3D Representations of Biomolecular Structures": User instructions

**Supplementary Information**

Jiyao Wang^1*^, Philippe Youkharibache^1,2^, Dachuan Zhang^1^, Christopher J. Lanczycki^1^, Renata C. Geer^1^, Thomas Madej^1,2^, Lon Phan^1^, Minghong Ward^1^, Shennan Lu^1^, Gabriele H. Marchler^1^, Yanli Wang^1^, Stephen H. Bryant^1^, Lewis Y. Geer^1^, Aron Marchler-Bauer^1^

^1^National Center for Biotechnology Information, National Library of Medicine, National Institutes of Health,
8600 Rockville Pike, Bethesda, MD, 20894, USA

^2^National Cancer Institute, National Institutes of Health, 8600 Rockville Pike, Bethesda, MD, 20894, USA

*

**1 Implementation**

**1.1 Third-party Libraries and Tools**

iCn3D Structure Viewer is a WebGL-based 3D viewer using Three.js and jQuery. Its source code is available at GitHub (<https://github.com/ncbi/icn3d>). To view 3D structures, no installation is required. iCn3D displays the 3D structures directly in a web browser. One example is shown in Figure 1, which can also be opened in a live browser window with the shortened URL https://icn3d.page.link/2rZWsy1LZmtTS3kBA. iCn3D has been developed based on the 3D styles of iview (Li, et al., 2014) and GLmol, the surface display of 3Dmol (Rego and Koes, 2015), and the sphere and cylinder display of NGL using the impostor technology (Rose, et al., 2018; Rose and Hildebrand, 2015).

**1.2 Data Types**

iCn3D reads 3D structure data from a variety of sources as shown in Table 2. The type of input structure identifier determines the file format that will be loaded into iCn3D for a given structure, and may slightly affect details of the display. The MMTF ID is preferable to PDB ID and mmCIF ID because MMTF data cover all 3D structures, are highly compressed, include DSSP-calculated secondary structures if the secondary structures are not provided originally, and include bond types such as double bond or triple bonds in the chemicals. The MMDB ID loads a structure in NCBI’s MMDB file format, which provides unique views not available in the other formats, such as the 2D interaction schematic display. iCn3D also reads local PDB files, mmCIF files, Mol2 files, SDF files, XYZ files, or files from a server via URLs.

**Table 2:** iCn3D reads 3D structure data from a variety of sources. For example, the URL to display the PDB ID 1TUP is formatted as the base URL “https://www.ncbi.nlm.nih.gov/Structure/icn3d/full.html?” plus the parameter “pdbid=1tup”.

| **Data Type** | **URL Parameter** |
| --- | --- |
| PDB ID | pdbid=1tup |
| mmCIF ID | mmcifid=1tup |
| MMTF ID (Bradley, et al., 2017) | mmtfid=1tup |
| NCBI MMDB ID | mmdbid=1tup |
| NCBI gi | gi=1310961 |
| NCBI VAST+ alignment | align=1HHO,4N7N |
| NCBI PubChem (Kim, et al., 2016) CID | cid=2244 |

Different views in iCn3D (such as the 2D interaction schematic) can be shown or hidden using URL parameters shown in Table 3. Users can also add commands in the URL directly using a parameter such as “&command=color+secondary+structure”.

**Table 3:** Different views in iCn3D can be shown or hidden using URL parameters. For example, the URL to show the 2D interaction schematic is formatted as the base URL “https://www.ncbi.nlm.nih.gov/Structure/icn3d/full.html?mmdbid=1tup” plus the parameter “&show2d=1”.

| **View** | **Parameter to Show** | **Parameter to Hide** |
| --- | --- | --- |
| 2D Interaction Schematic | &show2d=1 | &show2d=0 (default) |
| Biological Annotations | &showanno=1 | &showanno=0 (default) |
| Defined Sets | &showsets=1 | &showsets=0 (default) |
| Menu | &showmenu=1 (default) | &showmenu=0 |
| Command Window | &showcommand=1 (default) | &showcommand=0 |

**1.3 Selection Mode**

All operations in iCn3D apply to either all atoms, or to the current selection. By default, iCn3D uses the “All atoms” selection mode and applies the “style” and “color” options to all atoms. When users select part of the structure, the mode becomes “Selection”, and the mode switch button (next to the “Help” menu), as well as the “Style” and “Color” menus, become orange. The next operation applies only on the selected atoms. If users want to change style or color for all atoms, they can toggle the mode button back to “All atoms".

**1.4 Hydrogen Bonds in Chemical Binding**

Users can select “Chemical Binding” in the View menu to view hydrogen bonds between chemicals and proteins. A pair of atoms form a hydrogen bond if their elements are among nitrogen, oxygen, or fluorine and the distance between them is less or equal to 3.5 angstrom. The side chains of hydrogen donors or acceptors are shown in the Stick style. Hydrogen bonds are shown as dashed green lines. The view zooms in to all chemicals and their hydrogen-bonded residues. If users are interested in only one chemical, they can select the chemical by holding the “Alt” key and clicking on the chemical in the 3D structure, and then click “H-Bonds to Selection” in the View menu to show the hydrogen bonds with this chemical.

**1.5 Surface**

iCn3D can compute and display multiple surfaces for multiple selections. Three types of surfaces are available: Van der Waals Surface, Molecular Surface, and Solvent Accessible Surface (Xu and Zhang, 2009). By default, the surfaces are computed for selected atoms, without considering surrounding atoms. Users can select surface types containing “with Context” to show the surfaces considering the surrounding atoms. The color of a surface segment is determined by the color of the proximal atoms. Surfaces are generated with a maximum grid size of 180 angstroms. If the size of the structure is larger than the maximum grid size, a scale factor is used to fit the structure into this maximum grid size.

**1.6 Electron Density Map and EM Density Map**

Figure 4 shows both electron density map and B-factor (temperature factor) values for the PDB structure 3GVU. The 2Fo-Fc electron density map (contours in cyan) is contoured at 1.5 σ by clicking “Electron Density > 2Fo-Fc Map” in the Style menu. The structure is colored by B-factor values in the range of 0 (blue) -100 (red). The thickness of the B-factor tube is proportional to the B-factor value. The electron density map is consistent with the B-factor values: residues with large B-factor (white thick tubes) have poor electron density map.

iCn3D can show both 2Fo-Fc and Fo-Fc electron density maps for any subset of a crystal structure using the DSN6 file retrieved from RCSB. Users can choose a sigma value between 0 and 10 σ for the electron density maps. The default value is 1.5 σ for 2Fo-Fc map and 3.0 σ for Fo-Fc map. In the Fo-Fc map, the positive sigma values are colored in green and the negative sigma values are colored in red.

iCn3D can also show EM density map for any subset of an EM structure using the BinaryCIF file retrieved from the density server at PDBe (Sehnal, et al., 2017). The EM density values are linearly ranged from the minimum value to the maximum value. Users can choose the percentage from 0% (minimum) to 100% (maximum). The default percentage is 30%.

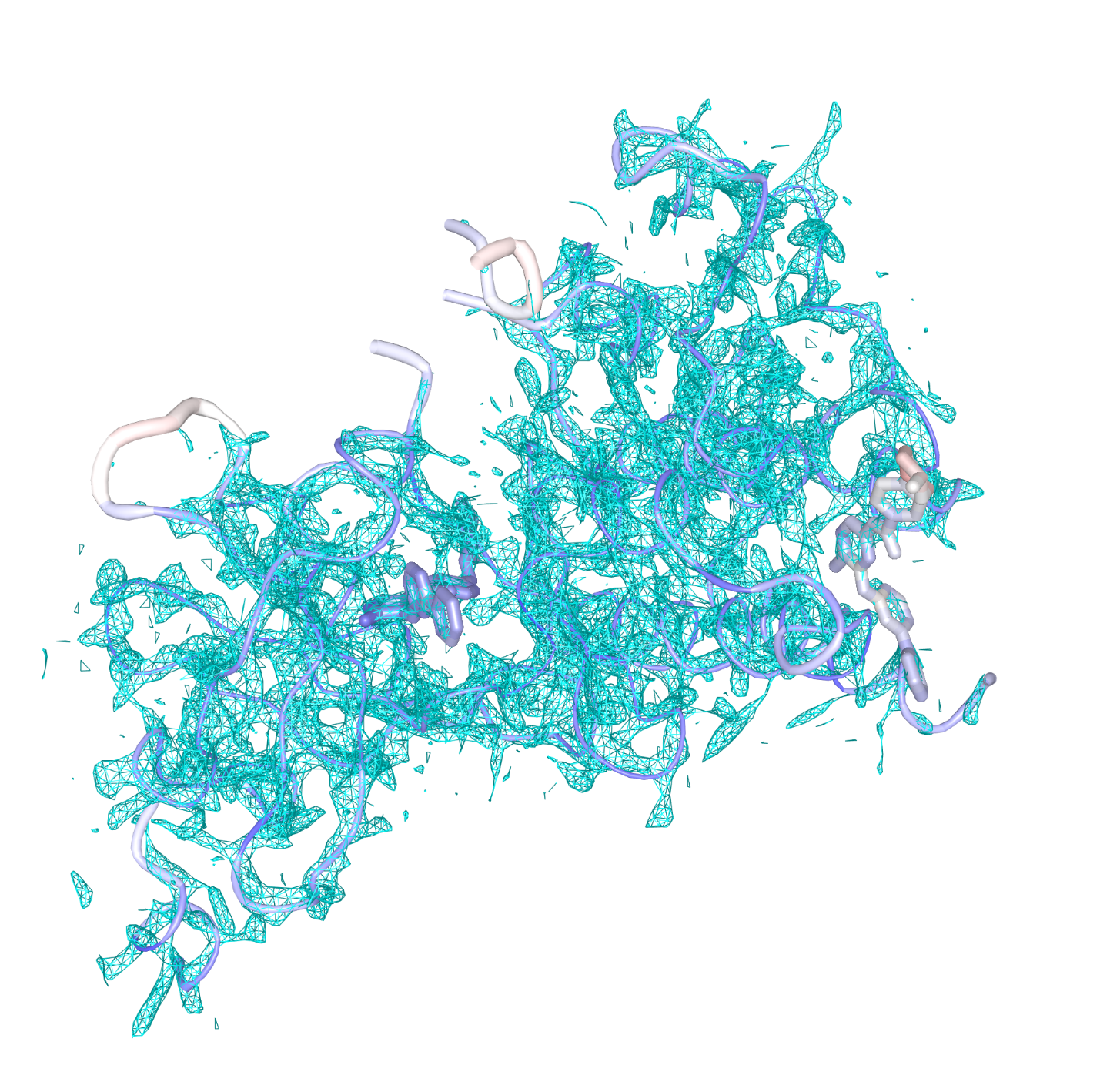

Figure 4. Electron density map and B-factor values of the PDB structure 3GVU. The 2Fo-Fc electron density map is contoured at 1.5 σ. The tube represents the asymmetric unit of the crystal structure. Its color changes from blue to red as the B-factor increases from 0 to 100. The tube is thicker when the B-factor value is larger. Share link: https://icn3d.page.link/cGcKMwbGFNRrzmYg8.

**1.7 Style**

Several commonly used representations such as Ribbon, Cylinder and Plate, C Alpha Trace, B-factor Tube, Stick, Sphere, etc. are available to display molecules.  In addition, iCn3D has a novel display style called “Schematic”, which is available for proteins, nucleotides, and chemicals. The schematic style shows one-letter residue abbreviations in the positions of the C-alpha atoms for proteins or the O3’ atoms for nucleotides. For chemicals, the schematic style renders bonds as sticks, and labels all non-carbon and non-hydrogen atoms with their element names.

The shiny surface display for 3D structures is achieved by using the material MeshPhongMaterial in Three.js. The arrows in the ribbon style of beta sheets are achieved by adding a triangle at the end of each strand. The triangle follows the curve of the strand, which are smoothed using the cubic spline algorithm.

If the structure is specified via a PDB ID or a PDB file, and the secondary structure assignments are missing, the DSSP method (Sicheri, et al.) is used to calculate the secondary structures.

**1.8 Color**

iCn3D offers many coloring schemes: “Spectrum”, “Secondary”, “Charge”, “Hydrophobic”, “Chain”, “Residue”, “Atom”, “B-factor”, “Unicolor”, and “Color Picker”. The “Charge” coloring scheme renders Arg and Lys residues in blue, and all nucleotides, Asp and Glu residues in red. The “Hydrophobic” coloring scheme renders Gly, Pro, Ala, Val, Leu, Ile, and Phe residues in green, and the remaining residues in grey. iCn3D also allows users to pick a color using “Color Picker” (https://github.com/tovic/color-picker).

**1.9 3D Printing**

iCn3D enables users to export STL or VRML files for 3D printing. An STL file can be printed in most 3D printers, but it has only one color. A VRML file can be used for 3D color printing. Some common 3D display styles such as “Ribbon” are suitable for 3D visualization but may require extra supporting structures/material in 3D printing. iCn3D adds default stabilizers if the export method is “STL with Stabilizers” or “VRML (Color, with Stabilizers)” in the File menu. The stabilizers for nucleotides are the hydrogen bonds. The stabilizers for proteins are connections made from every third residue to another residue that is within 4 angstroms. The stabilizers for chemicals and ions are connections from every 10th atoms in the chemicals or ions to another residue that is within 4 angstroms. Users can customize the size of the 3D objects by clicking “3D Printing > Set Thickness” in the File menu.

To build a printable 3D model, users can click “3D Printing > Add All Stabilizers” in the File menu to evaluate whether the 3D model appears stable. If not, they may click “Add One Stabilizer” to add extra stabilizers. Then users can click “STL” or “VRML (Color)” to export files. When users print a large biological assembly (e.g., a virus with 60 asymmetric units), each asymmetric unit is exported as a separate file. Users can then use external software to merge these files into a single file for 3D printing.

**1.10 Commands**

iCn3D has a defined command associated with each step. All commands are sequentially saved in memory. These commands enable users to share URL links (e.g., https://icn3d.page.link/2rZWsy1LZmtTS3kBA), save the visualization’s current state in a state file, undo and redo display changes, and directly modify the display using commands in the grey command window that appears beneath the 3D display.

The command corresponding to each operation is shown in that command window. Users can also find both the commands and the corresponding JavaScript functions by clicking “Web APIs > Commands” in the Help menu. On the Web APIs page, the “Menu”, “Command”, and “Method” columns list the operation name in the menus, the command name, and the JavaScript function, respectively.

**1.11 Share link**

One important feature of iCn3D is the facility to share a display, which includes both selections and rendering choices, with other users through a single URL. One example URL is https://icn3d.page.link/2rZWsy1LZmtTS3kBA (Figure 1), which is shortened using google Firebase Dynamic Links from the original URL: [https://www.ncbi.nlm.nih.gov/Structure/icn3d/full.html?mmdbid=1tup&showanno=1&show2d=1&showsets=1&command=view+annotations;+set+annotation+all;+set+view+detailed+view;+select+saved+atoms+1TUP_E+or+1TUP_F;+color+atom;+style+nucleotides+ball+and+stick;+select+.B:155;+select+zone+cutoff+4;+select+saved+atoms+1TUP_E+or+1TUP_F+and+sphere.B:R155-4A;+set+surface+Van+der+Waals+surface;+select+saved+atoms+1TUP_B+or+1TUP_E+or+1TUP_F;+show+selection;+add+track+|+chainid+1TUP_B+|+title+Custom+Key+Sites+|+text+82+R,+152+G,+155-156+RR,+180+R,+189+R;+select+.B:82,156,180,189;+color+blue;+add+residue+labels;+select+.B:152;+color+grey;+add+residue+labels;+select+.B:155;+color+blue;+add+label+R248+|+x+51.76+y+21.37+z+83.13+|+size+40+|+color+ffff00+|+background+cccccc+|+type+custom;+select+.B:82,152,155-156,180,189+|+name+keysites;+style+sidec+sphere|||{"factor":"1.1127","mouseChange":{"x":"0.065352","y":"0.032319"},"quaternion":{"_x":"0.10007","_y":"-0.22373","_z":"0.21026","_w":"0.94643"}}](https://www.ncbi.nlm.nih.gov/Structure/icn3d/full.html?mmdbid=1tup&showanno=1&show2d=1&showsets=1&command=view+annotations;+set+annotation+all;+set+view+detailed+view;+select+saved+atoms+1TUP_E+or+1TUP_F;+color+atom;+style+nucleotides+ball+and+stick;+select+.B:155;+select+zone+cutoff+4;+select+saved+atoms+1TUP_E+or+1TUP_F+and+sphere.B:R155-4A;+set+surface+Van+der+Waals+surface;+select+saved+atoms+1TUP_B+or+1TUP_E+or+1TUP_F;+show+selection;+add+track+|+chainid+1TUP_B+|+title+Custom+Key+Sites+|+text+82+R,+152+G,+155-156+RR,+180+R,+189+R;+select+.B:82,156,180,189;+color+blue;+add+residue+labels;+select+.B:152;+color+grey;+add+residue+labels;+select+.B:155;+color+blue;+add+label+R248+|+x+51.76+y+21.37+z+83.13+|+size+40+|+color+ffff00+|+background+cccccc+|+type+custom;+select+.B:82,152,155-156,180,189+|+name+keysites;+style+sidec+sphere|||%7b%22factor%22:%221.1127%22,%22mouseChange%22:%7b%22x%22:%220.065352%22,%22y%22:%220.032319%22%7d,%22quaternion%22:%7b%22_x%22:%220.10007%22,%22_y%22:%22-0.22373%22,%22_z%22:%220.21026%22,%22_w%22:%220.94643%22%7d%7d). The original URL includes the following commands, where “+” symbols are replaced by spaces. First, show sequences and annotations, 2D interaction schematic, and defined sets for the PDB structure 1TUP using the parameters “showanno=1”, “show2d=1”, and “showsets=1”. Second, show all annotations and view the sequence details using the commands “view annotations; set annotation all; set view detailed view;”. Third, color the nucleotide chains by atoms and display them with the style of ball and stick using the commands “select saved atoms 1TUP_E or 1TUP_F; color atom; style nucleotides ball and stick;”. Fourth, show the surface of nucleotides within 4 angstroms from Arg 248 in the chain B. When the input is MMDB ID, iCn3D uses NCBI residue numbers instead of PDB residue numbers. These two numbers are the same for most structures. But they are different by 93 for the PDB structure 1TUP since NCBI residue number always starts from 1. Thus, the corresponding NCBI residue number for Arg 248 is 155. We use the following commands to show the surface: “select .B:155; select zone cutoff 4; select saved atoms 1TUP_E or 1TUP_F and sphere.B:R155-4A; set surface Van der Waals surface;”. Fifth, display only the chains 1TUP_B, 1TUP_E, and 1TUP_F using the commands “select saved atoms 1TUP_B or 1TUP_E or 1TUP_F; show selection;”. Sixth, add a custom track with the key residues arginine 175, 248-249, 273 and 282, and glycine 245 using the commands “add track | chainid 1TUP_B | title Custom Key Sites | text 82 R, 152 G, 155-156 RR, 180 R, 189 R;”. Note the residue numbers used in the commands are 93 smaller than the “PDB Residue Numbers”. Seventh, color the key residues Arg in blue and Gly in grey and add residue labels using the commands “select .B:82,156,180,189; color blue; add residue labels; select .B:152; color grey; add residue labels; select .B:155; color blue; add label R248 | x 51.76 y 21.37 z 83.13 | size 40 | color ffff00 | background cccccc | type custom;”. Eighth, display the side chains of the key residues with the style of sphere using the commands “select .B:82,152,155-156,180,189 | name keysites; style sidec sphere;”. Finally, the orientation of the 3D display is set using the parameters after “|||”.

The Share Link above conveniently reproduces the custom display in Figure 1. This link can be shared with colleagues, inserted into presentations, or included in a publication, etc. It can be generated by clicking “Share Link” in the File menu. It is also available in the last line of each saved iCn3D PNG image, which is generated by clicking “Save Files > iCn3D PNG Image” in the File menu.  Images generated by clicking the right mouse button over the 3D display, or by printing the screen do not have the embedded link. Furthermore, this saved iCn3D PNG image can be used to reproduce the custom display by clicking “Open File > iCn3D PNG Image”. Thus, users can use these iCn3D PNG images to save their custom displays or reproduce these custom displays later.

**1.12 Selection on 3D**

To select the residue Arg with NCBI residue number 155, users can put the mouse over the residue in the groove of the structure’s 3D view (Figure 1A) to see the mouseover text “Arg155”, then hold the “Alt” key and click on the residue. The Ray Casting technology (https://threejs.org/docs/#api/core/Raycaster) is used to pick an atom in the structure. Once an atom is picked, the residue containing the atom is selected by default. Other options under “Select on 3D” in the Select menu include “Chain”, “Strand/Helix”, “Atom”. Once an atom is selected, users can use the up and down arrows to switch among different levels: atom, residue, strand/helix, chain, and the whole structure. Users can hold the “Ctrl” key to combine selections and hold the “Shift” key to select a range of residues. To toggle the highlight or remove the labels, users can click on “Show Toolbar” at the top-right corner and click on the button “Toggle Highlight” or “Remove Labels”.

**1.13 Selection on 2D**

To select chain B on the 2D interaction schematic (Figure 1B bottom), users can click on the node with the label B in the interaction schematic. Users can also click on the line that connects chain B with chain E. This operation selects the residues in chain B that interact with chain E. Two residues are considered as interacting residues if any of their non-hydrogen atoms are found within 4 angstroms of each other. The selected residue set is saved with a name “inter_B_E” in the list of “Defined Sets”. If only part of the chain is selected, a smaller node is shown inside the original node in the 2D interaction schematic. Selections on 2D interactions can be combined by holding the “Ctrl” key. Only the selected chains on 2D interactions show up in the “Sequences and Annotations” window.

**1.14 Selection on Defined Sets**

Users can also click on “1TUP_E” in the list of “Defined Sets” (Figure 1B top) to select the DNA chain E. Only the selected chain E will show up in the “Sequences and Annotations” window. Users can hold the “Ctrl” key to select multiple sets in “Defined Sets” and choose to display only these sets by clicking the button “View Only Selection” in the View menu. Users can also change the “union/or” of multiple sets to “intersection/and” or “exclusion/not” by clicking the “+” sign after “Set Operations” and clicking the button “Save Selection” in the window of “Defined Sets”. For example, if users click four sets ":1-10", ":11-20", ":5-15", and ":7-8" in the "Defined Sets", the command "saved atoms :1-10 or :11-20 or :5-15 or :7-8" will show up after the text “Select”. Users can modify the command to "saved atoms :1-10 or :11-20 and :5-15 not :7-8", which combines all residues 1-10 and 11-20 to get the residues 1-20, then intersects with the residues 5-15 to get the residues 5-15, then exclude the residues 7-8 to get the final residues 5-6 and 9-15.

**1.15 Selection on 1D**

In the 1D “Sequences and Annotations” window (Figures 1C and 1D), iCn3D provides annotations originating from various NCBI databases: SNPs (from dbSNP (Sherry, et al., 2001)), ClinVar (annotated SNPs from ClinVar (Landrum, et al., 2018)), Conserved Domains (from CDD (Marchler-Bauer, et al., 2017)), 3D Domains (from MMDB (Madej, et al., 2014)), Functional Sites (from CDD), and Interactions (from MMDB). Each annotation contains the title followed by the sequence. Users can click on the title to select all sequences on this track. To select multiple tracks, click the titles while holding the “Ctrl” key. To select a few residues in the sequence, users can simply use mouse to drag on the sequence. Each additional drag adds more residues to the selection. To deselect a residue, drag on the residue again. To show the sequences and annotations of all chains, click the button “Show All Chains” in this window. Users can click on the buttons “< ClinVar” and “ClinVar >” to browse through ClinVar (clinically relevant variants) annotation, which will be highlighted and labeled on the 3D display. Users can also mouse over the residues on the ClinVar track to see details about each variant.

SNPs and ClinVar tracks show the pathogenic variants in purple, other ClinVar clinically significant variants in green, and variants that are not in ClinVar but are in dbSNP in black (Figure 1D). (dbSNP annotates variants on RefSeq (O'Leary, et al., 2016; Rajput, et al., 2019) protein sequences as part of its regular build release process. The mapping of dbSNP variants to PDB structures was done by using the annotated RefSeq protein sequences to BLAST (Altschul, et al., 1997; Altschul, et al., 2005) search against PDB database. The BLAST alignments were filtered to include only the top PDB protein hits with E-value less than 0.0001 and were used to project the variants from RefSeq proteins to the corresponding positions on the PDB proteins.) Conserved Domains show the location of conserved domain footprints as pre-computed by CDD annotation services. 3D Domains are domain models based on 3D structures and reflect compact structural units within a protein that have been identified using purely geometric criteria. Functional Sites are obtained from pre-computed CDD annotation and are defined as sets of residues that are associated with molecular function and presumed conserved in a domain family, such as binding sites, or active/catalytic sites. Interactions shows the residues in one molecule/chain that have contact with another molecule; the interactions are calculated on-the-fly, or pre-calculated at NCBI by identifying non-hydrogen atoms that are found within 4 angstroms of each other. They can also be selected from the 2D interaction schematic by clicking on the line that connects two molecules.

**1.16 Search Sequences and Advanced Selection**

iCn3D features a search button at the top-right corner. Users can search a segment of sequence in the one-letter format (e.g., “GXXRR”). Matched residues will be highlighted in 3D structure, 2D interactions, the list of Defined Sets, and 1D Sequences and Annotations. The matched residues are also saved as a defined set with the name such as “:GXXRR”. The symbol “:” means residues and “X” is a wildcard character to represent any single residue. The search shares the same specification as the “Advanced” option in the Select menu, except that it is not necessary to include the “:” symbol when using the “Search Seq.” box to find residues within sequences.

The “Advanced” selection is a powerful tool to build custom selections. Users can input a selection command with a simple specification and save the selection with a name. For example, in the selection “$1TUP.A,B:5-10,R,ions@CA,C”, the term “$1TUP” uses the prefix “$” to indicate a list of comma-separated structures; “.A,B” uses the prefix “.” to indicate macromolecules/chains; “:5-10,R,ions” uses the prefix “:” to indicate individual residues. Residues can be specified via residue numbers or ranges of residue numbers (e.g., “5-10”), one-letter residue codes (e.g., “R”), or via predefined names: “proteins”, “nucleotides”, “chemicals”, “ions”, and “water”. The prefix “@” is used to specify individual atoms. For example, “@CA,C” instructs iCn3D to select the alpha carbon and carbon atoms in the specified regions of the structure. Partial specifications are allowed, e.g., the selection “:5-10” selects residues with the residue numbers in the range of 5-10 in all structures and chains.

**1.17 Custom Tracks**

Users can also add custom tracks in the Sequences and Annotations window to complement or supplement existing annotations. To add custom tracks, click the button “Add Track” in each chain section. Currently there are five ways to add custom tracks. First, the current selection can be shown as a track. Users can make any selection and show the selection as a track. Second, users can input a protein sequence in the FASTA format. The sequence is aligned with the sequence of the current chain using the BLAST algorithm. If a match is confirmed, the matched part of the input sequence will be shown as a custom track. Otherwise, an error message “cannot be aligned” will be displayed. Third, users can input a BED file to show multiple tracks. The BED file can include color information. Fourth, users can add any custom text as a track (e.g., “Custom Key Sites” in Figure 1D). To save space, the input is a list of comma-separated position and text pairs. For example, "152 G, 155-156 RR" defines a character "G" at the position 152 and two continuous characters "RR" at positions from 155 to 156. The starting position is 1. Fifth, users can input NCBI protein accessions to add a track. The sequence of the protein will be aligned with the sequence of the current chain using BLAST, and the matched sequence will be shown as a custom track.

**1.18 Performance**

iCn3D displays large structures in a few seconds. For example, it can display the complex structures of the large ribosomal subunit of Haloarcula marismortui (<https://www.ncbi.nlm.nih.gov/Structure/icn3d/full.html?mmtfid=1ffk>) and of a virus capsid (<https://www.ncbi.nlm.nih.gov/Structure/icn3d/full.html?mmtfid=2bbv>) in 2-3 seconds, comparable to other 3D viewers such as NGL (Rose, et al., 2018; Rose and Hildebrand, 2015) and LiteMol (Sehnal, et al., 2017). These structures contain 64,281 and 469,020 atoms, respectively.

Several technologies are used to speed up the display. First, iCn3D uses the impostor technology (Rose, et al., 2018; Rose and Hildebrand, 2015) to render spheres and cylinders directly with shaders when the WebGL extension on fragment depth value is available in the browser (e.g., Chrome, Safari, Firefox, and Microsoft Edge, but not Internet Explorer). Since spheres and cylinders are used a lot, the impostor technology speeds up the rendering significantly. Second, iCn3D uses Instanced Buffer Geometry in Three.js to render biological assemblies. Each biological assembly consists of many copies of an asymmetric unit. The Instanced Buffer Geometry technology speeds up the display by creating the geometry of the asymmetric unit only once, transforming it according to the symmetry matrices to generate translated/rotated copies, and rendering the transformed copies. Third, iCn3D accepts the highly compressed binary MMTF data format. This greatly speeds up the loading of 3D coordinates.

**1.19 Embed iCn3D with iframe or JavaScript libraries**

iCn3D can be embedded in a web page by including the URL in HTML iframe, e.g. <iframe src="https://www.ncbi.nlm.nih.gov/Structure/icn3d/full.html?mmdbid=1tup&showanno=1&show2d=1&showsets=1&width=800&height=450&showmenu=1&showcommand=1&rotate=right" width="900" height="600" style="border:none"></iframe>. This method always shows the most recent version of iCn3D.

To embed iCn3D with JavaScript libraries, the following libraries need to be included: jQuery, jQuery UI, Three.js, and iCn3D library. An html div tag to hold the 3D viewer is added: “<div id=’icn3dwrap’></div>”. The iCn3D widget is initialized with the custom defined parameter “cfg”: “var icn3dui = new iCn3DUI(cfg); icn3dui.show3DStructure();”. Multiple iCn3D widgets can be embedded in a single page. Users can choose to show the most recent version of iCn3D, or a locked version of iCn3D. To show the most recent version, use the library files without the version postfix as shown in the iCn3D Web API page (<https://www.ncbi.nlm.nih.gov/Structure/icn3d/icn3d.html#HowToUse>). To show a locked version, use the library files with the version postfix as shown in the source code of iCn3D page (<https://www.ncbi.nlm.nih.gov/Structure/icn3d/full.html?mmdbid=1tup>). If the input is provided as an MMDB ID, both library files and backend cgis are versioned so that the 3D display will be stable.

**2 Web Applications**

(a)	(b)

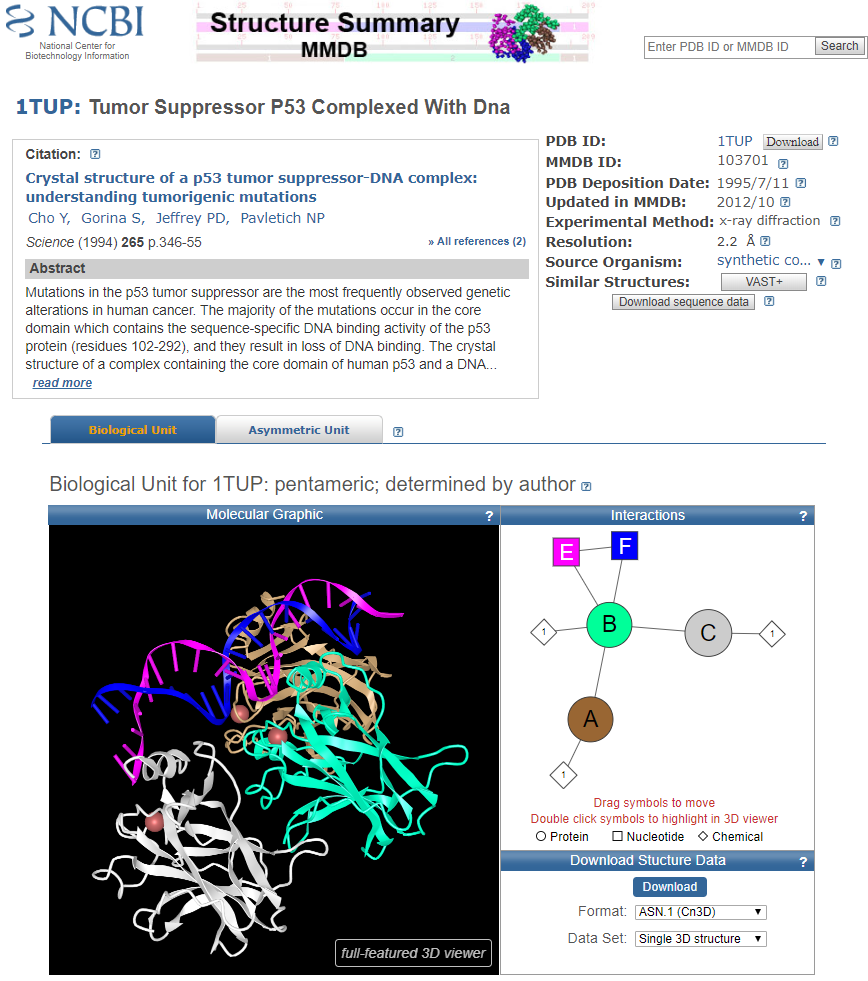

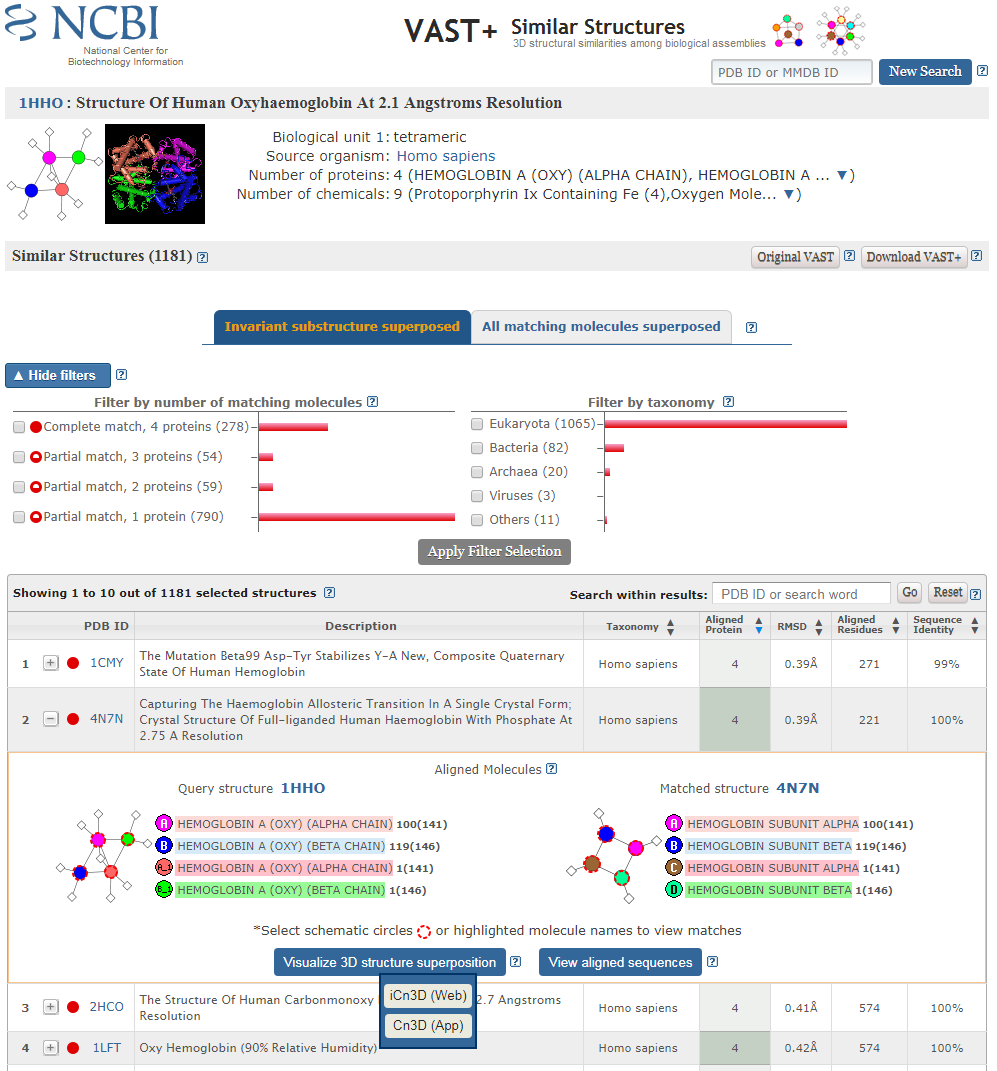

**Figure 5:** (a) MMDB page showing the PDB structure 1TUP. (Link: <https://www.ncbi.nlm.nih.gov/Structure/pdb/1TUP>) (b) VAST+ page showing the similar structures of the PDB structure 1HHO. (Link: <https://www.ncbi.nlm.nih.gov/Structure/vastplus/vastplus.cgi?uid=1HHO>)

As shown in Figure 5a, iCn3D has been implemented as the default viewing option in the structure summary pages of NCBI’s structure database MMDB (e.g., <https://www.ncbi.nlm.nih.gov/Structure/pdb/1TUP>). The interactive 3D displays can be rendered without requiring users to install a stand-alone application, such as the Cn3D viewer (Wang, et al., 2000). The structure summary page shows a static molecular graphic image by default. Users can click the button “3D view” at the bottom-left corner of the image to view the structure using iCn3D or click the button “full-featured 3D viewer” at the bottom-right corner to launch iCn3D in a separate window.

As shown in Figure 5b, iCn3D has also been implemented as a viewer in the VAST+ resource pages, which summarize the results of 3D comparisons between similar macromolecular complexes, including superpositions and corresponding alignments, e.g., <https://www.ncbi.nlm.nih.gov/Structure/vastplus/vastplus.cgi?uid=1HHO>. To view structure superpositions, users need to select one of the aligned structures and expand the corresponding table row to show details about the comparison. In the resulting panel, click on the button “Visualize 3D structure superposition”, and choose “iCn3D (Web)” in the popup to launch iCn3D.
